# Production of Glycolic acid through Whole-Cell Bioconversion from PET Monomer Ethylene Glycol Using Engineered *Corynebacterium glutamicum*

**DOI:** 10.1101/2025.07.06.663337

**Authors:** Mohammad Rifqi Ghiffary, Fong Tian Wong, Yee Hwee Lim

**Affiliations:** Institute of Sustainability for Chemicals, Energy and Environment (ISCE2), Agency for Science, Technology and Research (A*STAR), 1 Pesek Road, Jurong Island, Singapore 627833, Republic of Singapore; Singapore Integrative Biosystems and Engineering Research (SIBER), Agency for Science, Technology and Research (A*STAR), 20 Biopolis Way, #08-01 Centros, Singapore 138668, Republic of Singapore; Molecular Engineering Lab, Institute of Molecular and Cell Biology (IMCB), Agency for Science, Technology and Research (A*STAR), 61 Biopolis Drive, #07-06, Proteos, Singapore, 138673, Republic of Singapore; Synthetic Biology Translational Research Program, Yong Loo Lin School of Medicine, National University of Singapore, 10 Medical Drive, Singapore, 117597, Republic of Singapore

**Author notes:** Corresponding authors: Email addresses.

**Keywords:** *Corynebacterium glutamicum*, metabolic engineering, biotransformation, ethylene glycol, glycolic acid

## Abstract

In the last decade, the global warming and plastic pollution issue have driven research on developing a more sustainable platform for chemicals production from alternative feedstocks. Ethylene glycol (EG), a monomer of polyethylene terephthalate (PET) plastic, has a potential to become a renewable substrate for microbial production of value-added chemicals. This study presents a biotransformation platform using *Corynebacterium glutamicum* to produce glycolic acid (GA) from EG. *C. glutamicum* was engineered to express a heterologous EG oxidation pathway. Subsequent promoter engineering yielded strain FA4, producing 10.6 g/L GA from EG in 48 h. Implementation of a two-stage biotransformation strategy using resting cells further enhanced the GA production, reaching a cumulative GA titer of 98.8 g/L after a 72-h production. Finally, applying this platform to a simulated EG mixture from PET-degradation achieved a cumulative GA titer of 67.3 g/L over 72 h, highlighting the potential for valorizing plastic waste through this biotransformation platform. These findings establish *C. glutamicum* as an efficient biotransformation chassis for sustainable GA production from EG and offer a promising route for PET waste valorization into value-added chemicals.

## 1. Introduction

In recent years, the growing plastic pollution crisis has prompted the development of innovative recycling technologies to mitigate environmental impact. Polyethylene terephthalate (PET) for example, as one of the world’s widely utilized plastics, has garnered attention due to its potential for recycling into constituent monomers, including terephthalic acid (TPA) and ethylene glycol (EG) (Ragaert et al., 2017). While substantial development of upcycling technologies for PET monomers have been made, the focus has predominantly centred on recycling TPA but leaving EG underutilized (Pandit et al., 2021). This disparity comes from the fact that recycling the EG fraction of PET hydrolysate is not economically attractive relative to its high energy requirements for its distillation. Alternatively, the EG solution generated from PET plastic degradation processes could directly serve as carbon substrate for biotechnological conversions which can effectively by-pass the EG purification challenge and serve as innovative method for EG upcycling (Balola et al., 2024). Additionally, utilizing EG as carbon source can promote diversification of alternative feedstocks in industrial biotechnology, avoid competition with other important industries (e.g. foods), and ultimately create a circular bioeconomy.

Building upon the promise of waste-derived EG as a sustainable carbon source, we were particularly interested in upcycling EG into glycolic acid (GA) to significantly elevate the economic and environmental value proposition of this feedstock. GA is an α-hydroxy acid with a rapidly expanding global market, driven by its versatile applications in industries such as cosmetics (as an exfoliant and anti-aging agent), textiles (for dyeing and finishing), pharmaceuticals (as a precursor for drug synthesis and biodegradable sutures), and as a monomer for environmentally friendly biopolymers like polyglycolic acid (Pandya et al., 2025; Sharad, 2013; Shrestha et al., 2024). Traditionally, GA is synthesized through fossil-fuel-dependent chemical routes, which are often energy-intensive and can involve hazardous reagents (Balola et al., 2024; Lee et al., 2023). Therefore, developing a sustainable, bio-based production method for GA, particularly from waste-derived EG, offers substantial environmental and economic advantages by transforming a lower-value, impure waste stream into a higher value, readily marketable chemical (Fig. 1A).

**Fig. 1.**
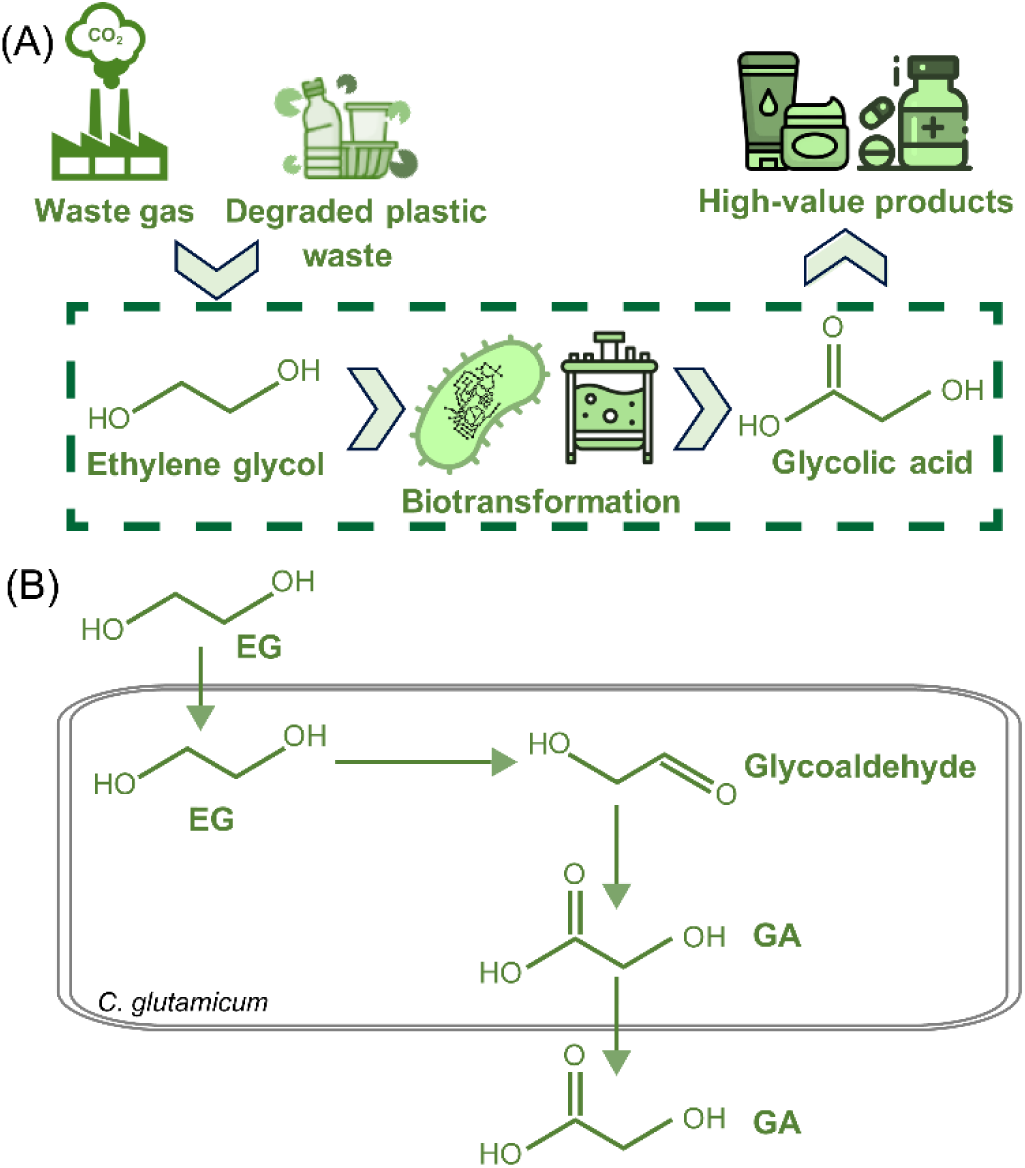
Overview for biotransformation platform for glycolic acid (GA) production from ethylene glycol (EG). (A) Waste gas (e.g., CO_2_) and degraded PET-plastic waste can be the potential source of EG as a sustainable feedstock for biotransformation to GA using engineered microbial cells. (B) Simplified metabolic pathway for the conversion of EG to GA. The pathway involves key oxidation steps, with EG initially oxidized to glycolaldehyde by propanediol or EG oxidoreductase, which is then further oxidized to the final product, GA by glycolaldehyde dehydrogenase.

To date, several studies have reported the natural capabilities of microbial species to assimilate EG as carbon and energy source. Notable examples include *Flavobacterium* sp. (Willetts, 1979), *Clostridium glycolicum* (Gaston and Stadtman, 1963), *Gluconobacter oxydans* (Santos et al., 2024; Zhang et al., 2015), *Yarrowia lipolytica* (Carniel et al., 2023), and *Rhodococcus jostii* (Roccor et al., 2024). Microbial EG assimilation can be divided into two different routes, aerobic and anaerobic, based on its dependence on oxygen. Aerobic ethylene glycol (EG) assimilation typically follows a conserved pathway involving its stepwise oxidation to glyoxylate. Anaerobic EG catabolism uses diol dehydratase to dehydrate EG into acetaldehyde (Shimizu and Inui, 2024). While the enzymes and cofactors responsible for the initial EG oxidation differ among microbial species, the resulting glyoxylate is subsequently metabolized through either the glycerate pathway or the β-hydroxyaspartate cycle. Of note, many of these microorganisms are not considered ideal industrial hosts due to limitations in genetic accessibility, robustness, or scalability even though they demonstrate a foundational ability to metabolize EG. Recognizing these limitations, significant efforts have been directed towards engineering well-established model microorganisms like *Escherichia coli* and *Pseudomonas putida* for improved EG assimilation. Early work in *E. coli* K-12 MG1655 involved expressing *fucO* (encoding for propanediol oxidoreductase) and *aldA* (encoding for glycolaldehyde dehydrogenase), enabling initial growth on EG when 4 g/L glycerol was also supplemented (Szappanos et al., 2016). However, engineering *E. coli* for high EG assimilation and subsequent product formation has faced hurdles, primarily due to the toxicity of intermediate metabolites like glycolaldehyde and the inhibitory effects of EG and GA at elevated concentrations (Panda et al., 2021; Yan et al., 2024). Similarly, while *P. putida* demonstrates remarkable metabolic versatility, engineering it for specific EG bioconversion pathways has required extensive pathway balancing and engineering strategies to circumvent native metabolic bottlenecks and minimize substrate/product inhibition, often yielding moderate titers that still require further optimization for industrial viability (Franden et al., 2018; Mückschel et al., 2012).

One other well-established biotransformation chassis is *Corynebacterium glutamicum*, which has long been employed in the industrial production of amino acids and other biochemicals, due to its robust metabolism, abundant of genetic toolboxes availability, and high tolerance to diverse substrates (Becker et al., 2018; Cho et al., 2023). Recent advancements have extended its substrate range to include next-generation feedstocks such as methanol (Hennig et al., 2020), ethanol (Arndt et al., 2008), formate (Li et al., 2024), and acetate (Nickel et al., 2010). Interestingly, despite its potential, *C. glutamicum* has no known pathway to utilize EG directly as a primary carbon source, and to the best of our knowledge, there has been no studies on engineering *C. glutamicum* for EG assimilation.

Herein, we explored the feasibility of leveraging *C. glutamicum* as a biotransformation chassis to convert EG into GA (Fig. 1B). We first investigated the tolerance or toxicity limit of wild-type *C. glutamicum* to high concentrations of EG and GA. By testing different heterologous gene combinations from various species, we constructed a heterologous EG oxidation pathway in *C. glutamicum* and optimized its metabolic flux through promoter sequence engineering. Using a two-stage biotransformation strategy utilizing repetitive resting cells to overcome titers limitations, we demonstrated a high cumulative GA titer of nearly 100 g/L in 72 h production. Moreover, we further demonstrate the feasibility of a simulated EG-derived mixture from a post PET degradation as a feedstock for GA production, paving the way for upgrading PET-derived EG into value-added chemicals.

## 2. Materials and methods

### 2.1 Bacterial strains and culture conditions

All bacterial strains and plasmids employed in this study are detailed in Table 1, with primer sequences provided in Supplementary Table S1. *Escherichia coli* DH10B served as the host for routine cloning procedures. *Corynebacterium glutamicum* JCM 1318 (also catalogued as ATCC 13032) was obtained from the RIKEN BioResource Center through the National BioResource Project (MEXT, Japan). *E. coli* strains were cultivated in Luria-Bertani (LB) medium (10 g/L of NaCl, 10 g/L of tryptone, and 5 g/L of yeast extract) at 37 °C with agitation at 200 rpm, or on LB agar plates (1.5% w/v). *C. glutamicum* was maintained in brain heart infusion sorbitol (BHIS) medium (37 g/L of brain heart infusion and 91 g/L of D-sorbitol) at 200 rpm and 30 °C, or on BHIS plates (1.5%, w/v agar) for routine cultures. For glycolic acid (GA) production assays, *C. glutamicum* cultures were grown in CGXII minimal medium consisting of 40 g/L glucose, 20 g/L (NH_4_)_2_SO_4_, 5 g/L urea, 1 g/L KH_2_PO_4_, 1 g/L K_2_HPO_4_, 42 g/L MOPS, 0.25 g/L MgSO_4_·7H_2_O, 0.03 g/L protocatechuic acid, 5 mL/L trace metal solution, 200 µg/L biotin, and 500 μg/L thiamine-HCl. The pH was adjusted to 8.0 with NaOH. The trace metal solution was prepared as previously described (Luo et al., 2019), comprising 2.6 g/L CaCl_2_⋅2H_2_O, 2 g/L FeSO_4_⋅7H_2_O, 2.8 g/L MnSO_4_⋅5H_2_O, 0.2 g/L ZnSO_4_⋅7H_2_O, 0.06 g/L CuSO_4_⋅5H_2_O, 4 mg/L NiCl_2_⋅6H_2_O, and 10 mL/L HCl. Cultures were incubated in 250 mL baffled flasks containing 25 mL of modified CGXII medium at 30 °C and 200 rpm for 48 h. Where appropriate, kanamycin (50 µg/mL) and IPTG (1 mM) were added to the medium.

**Table 1.**
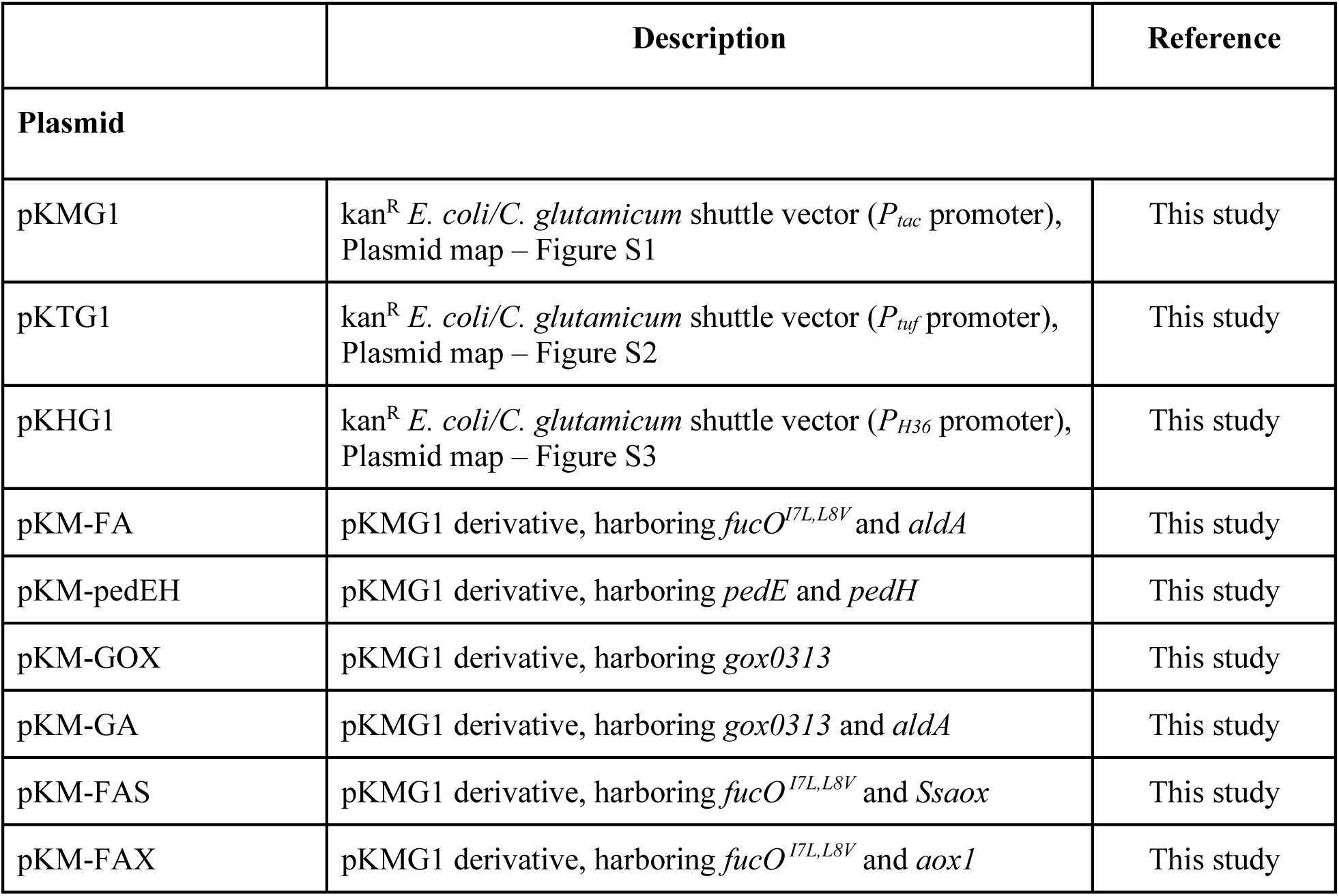
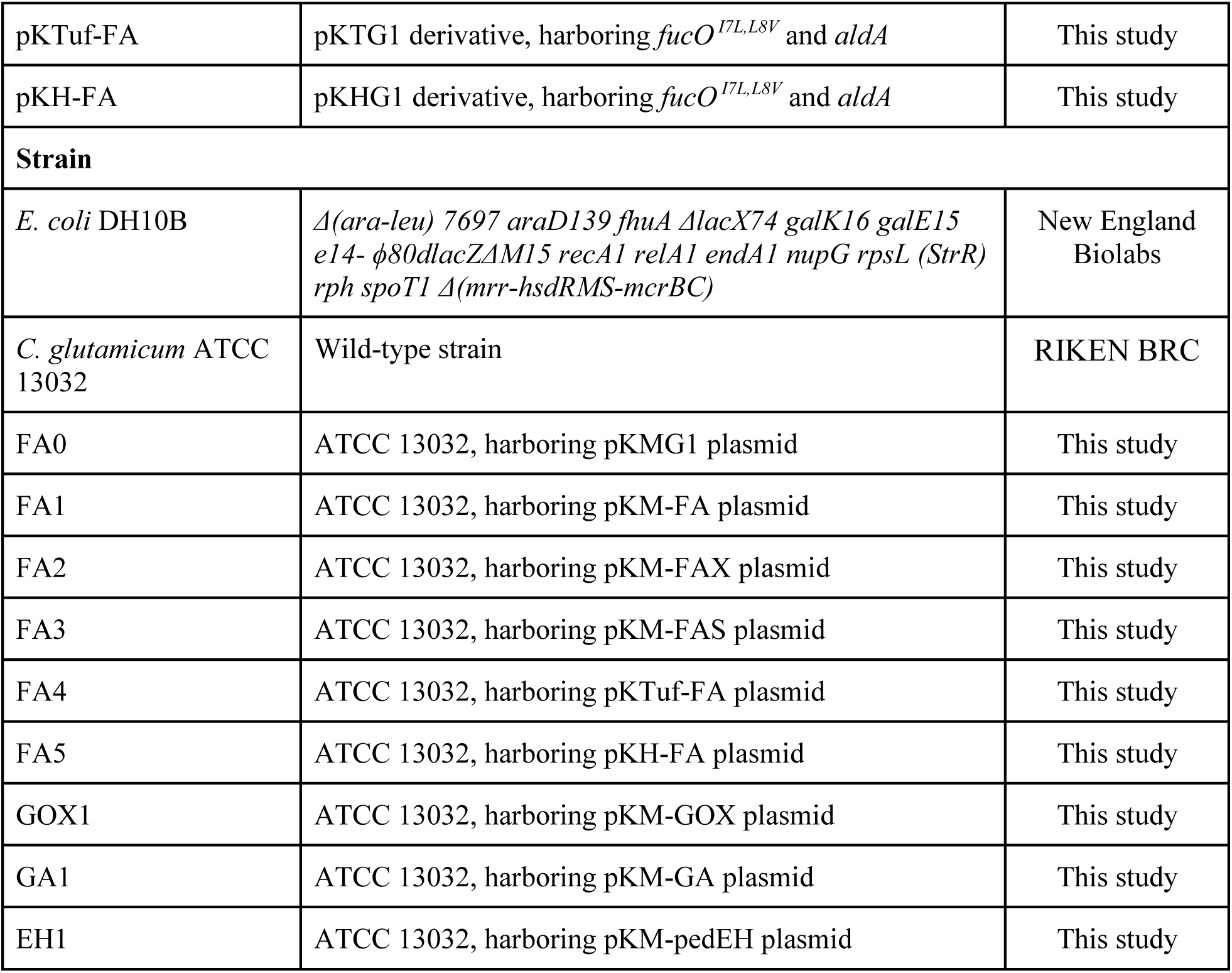
Plasmids and strains list.

### 2.2 Growth assays

Bacterial cell growth profiles in the presence of varying concentrations of EG and GA were monitored using a Varioskan LUX Multimode Microplate Reader (Thermo Fisher Scientific, USA). Overnight cultures of wild-type *C. glutamicum* were inoculated into 96-well microtiter plates containing 100 µL of CGXII minimal medium supplemented with 40 g/L glucose (as control, unless specified) and increasing concentrations of EG (0-100 g/L) or GA (0-100 g/L). Each well was inoculated to an initial optical density at 600 nm (OD_600_) of approximately 0.05. The plates were incubated at 30°C with continuous orbital shaking (480 rpm), and the OD_600_ of each well was measured automatically every 30 minutes for a total duration of 48 hours.

### 2.3 Cloning and expression

Molecular cloning procedures were performed using established protocols, including Gibson assembly and standard restriction-ligation methods (Gibson et al., 2009; Green et al., 2012). Codon-optimized synthetic gene sequences are listed in Supplementary Table S2. All recombinant plasmids were confirmed by Sanger sequencing. Transformation of plasmids into *C. glutamicum* was carried out following the method described by van der Rest et al. (1999).

### 2.4 Two-stage biotransformation

*C. glutamicum* cells were initially cultivated in CGXII minimal medium supplemented with 40 g/L glucose under the same condition as described above. Following the growth phase, the cells were harvested by centrifugation at 6000 rpm for 5 mins. The supernatant was discarded, and the cell pellet was resuspended in fresh CGXII minimal medium containing 40 g/L EG. The biotransformation was carried out in shake flasks at 30°C with shaking at 200 rpm for 24 hours. After each 24-hour cycle, samples were collected to measure biomass (OD_600_) and the concentrations of EG, GA, followed by cells harvesting and resuspension in fresh CGXII minimal medium containing 40 g/L EG for the next 24-hour biotransformation cycle. This process was repeated every 24 hours for a total of 3 days (72 hours of cumulative production time). For the biotransformation experiments using a mock EG mixture solution, the resting cells were prepared using the same procedure described above. The mock EG mixture solution was prepared in CGXII minimal medium (without glucose) containing 40 g/L EG, 2.5 g/L terephthalic acid (TPA), and 1 g/L polyethylene terephthalate (PET) with the initial pH solution of 8.0. The biotransformation in this mock solution was carried out under the same conditions and procedure as above.

### 2.5 Quantification of metabolites

For metabolite analysis, culture samples were centrifuged at 13,000 × g for 1 min. The supernatant was subsequently filtered through a 0.2 µm cellulose acetate membrane and diluted with deionized water up to appropriate concentration. Quantification of glucose, EG, and GA concentrations were calibrated and measured using HPLC on an Agilent 1200 series system (Agilent USA, Santa Clara, CA), equipped with a Phenomenex Rezex column (Phenomenex, Torrance, CA), and maintained at 60°C. A mobile phase of 0.005 N sulfuric acid was used at a flow rate of 0.6 mL/min and compound detection was done using a refractive index detector (RID).

## 3. Results and discussion

### 3.1 Tolerance of C. glutamicum to EG and GA

The potential of EG as a promising next-generation feedstock for biomanufacturing of chemicals using metabolic engineering is a relatively new area of exploration. Yet, the utilization of well-established industrial microorganisms, such as *Corynebacterium glutamicum* as a bioconversion chassis for this purpose has received limited attention. A previous work has indicated that both EG and GA can disrupt membrane integrity and metabolic homeostasis in microorganisms at elevated concentration (Yan et al., 2024). Similarly, glycolaldehyde, an intermediate molecule in the EG assimilation pathway, can also be detrimental to cells at higher levels (Yan et al., 2024). Therefore, we examined the impact of EG and GA on the growth of the wild-type *C. glutamicum* to evaluate its robustness as bioconversion chassis for GA production from EG. Cells were grown in CGXII minimal media with elevated concentrations of EG and GA and growth profiles (OD_600_) were measured every 30 mins for 48 h (see Materials and Method). Our results demonstrated that wild-type *C. glutamicum* could not grow in minimal media using EG or GA as the sole carbon source, suggesting that the organism does not harbour the requisite metabolite pathways to assimilate EG or GA directly (Fig. 2). However, when glucose was supplemented as the primary carbon source, *C. glutamicum* could grow and maintained significant growth even at high concentration of EG and GA. Interestingly, final biomass (OD_600_) of *C. glutamicum* is higher by 18 and 23% compared to control condition when 5 and 10 g/L EG were supplemented, respectively. The cell growth rate remained relatively high (0.067 h^-1^) at lower EG concentrations (up to 25 g/L), only slightly reduced from the control (0.077 h^-1^, Fig. 2D). As EG concentration further increased, the growth rate declined more sharply, reaching 0.047 h^-1^ at 50 g/L EG, 0.029 h^-1^ at 75 g/L EG, and 0.015 h^-1^ at 100 g/L EG. Despite this reduction, visible cell growth was still observed even at 100 g/L EG (Fig. 2D). A similar trend was observed for GA where the growth rates were minimally affected up to 25 g/L GA, albeit with longer lag phase (Fig. 2C). However, the *C. glutamicum* growth rate began to decrease more notably at 50 g/L GA and even inhibited the growth completely above this concentration (Fig. 2C). Comparatively, it has been reported that *E. coli* growth is inhibited in media containing more than 20 g/L EG due to the excessive accumulation of GA (around 4 g/L) as a major byproduct that can inhibit the activity of glyoxylate oxidase, thereby disrupting glyoxylate metabolism and ultimately impeding energy supply (Carniel et al., 2023; Franden et al., 2018). Thus, this demonstrates that *C. glutamicum* could be a more superior chassis for EG-to-GA conversion with significantly higher tolerance to both EG and GA compared to other model bacterial chassis.

**Fig. 2.**
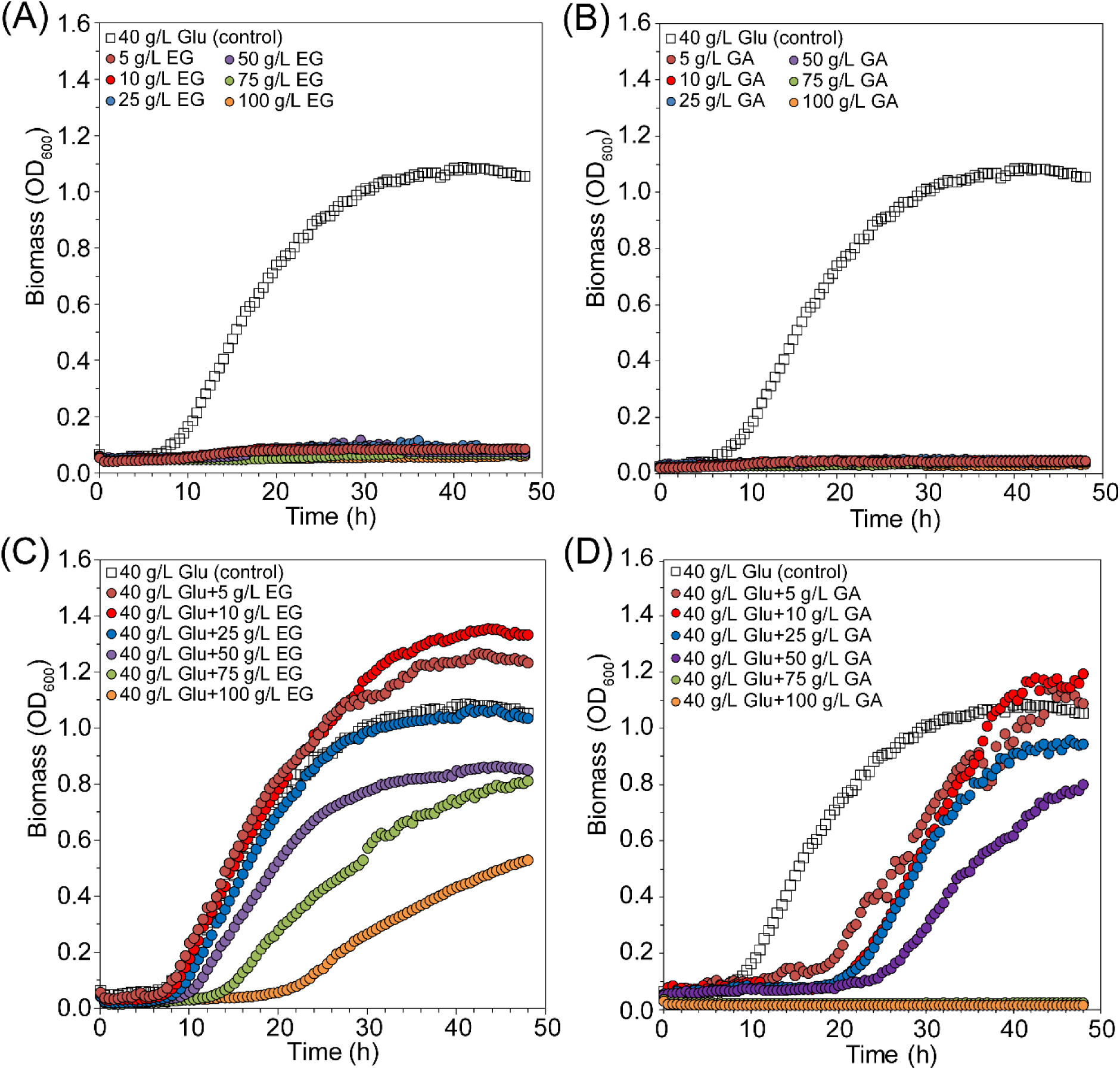
Tolerance test of *C. glutamicum* in elevated concentration of EG and GA. (A) Growth profile of *C. glutamicum* grown in CGXII minimal media containing 40 g/L glucose or various concentration of EG as sole carbon source. (B) Growth profile of *C. glutamicum* grown in CGXII minimal media containing 40 g/L glucose or various concentration of GA as sole carbon source. (C) Growth profile of *C. glutamicum* grown in CGXII minimal media supplemented with 40 g/L glucose and various concentrations of EG. (D) Growth profile of *C. glutamicum* grown in CGXII minimal media supplemented with 40 g/L glucose and various concentrations of GA. Results are given as the average of n = 3.

### 3.2 Construction of GA biosynthetic pathway from EG in C. glutamicum

The EG oxidation pathway has been recently elucidated in several bacteria, including *E. coli*, *Pseudomonas*, *Gluconobacter*, and *Flavobacterium* species. In this pathway, EG is initially converted to glycolaldehyde by a propanediol or EG oxidoreductase, or other alcohol dehydrogenases. Subsequently, glycolaldehyde is converted to GA by glycolaldehyde dehydrogenase (Balola et al., 2024). To the best of our knowledge, no EG oxidation pathway has been curated in *C. glutamicum* yet. Therefore, to construct *C. glutamicum* strains capable of converting EG to GA, several candidates genes were selected and evaluated, including *fucO*^I7L/L8V^ (propanediol oxidoreductase from *E. coli*), *gox0313* (alcohol dehydrogenase from *Gluconobacter oxydans*), *aldA* (glycolaldehyde dehydrogenase from *E. coli*), and *pedEH* (alcohol dehydrogenases from *P. putida*).

The *tac* promoter (P_tac_), which is inducible by IPTG, was initially employed to regulate gene expression and prevent excessive accumulation of glycolaldehyde, an intermediate product which can be detrimental to the cells. Subsequently, plasmids harboring *fucO*^I7L/L8V^ – *aldA* (pKM-FA), *gox0313* (pKM-GOX), *gox0313*-*aldA* (pKM-GA), and *pedE-pedH* (pKM-pedEH) were constructed and individually transformed into *C. glutamicum*, resulting in FA1, GOX1, GA1, and EH1 strains, respectively. The resulting transformants were then cultivated in shake flasks using CGXII minimal media supplemented with 10 g/L EG and 40 g/L glucose for 48 h. Compared to FA0 strain harboring the empty plasmid of pKMG1, all strains expressing heterologous genes showed significant EG assimilation, with the FA1 achieving the highest GA titer of 6 g/L (Fig. 3). Interestingly, the final biomass (OD_600_) of FA1 strains was significantly higher than that of the other strains, which might be attributed to the ability of FucO to perform inverse reduction reaction, thereby hindering the accumulation of toxic glycolaldehyde (Wagner et al., 2023). Nonetheless, while maintaining robust cell growth (OD_600_), our FA1 strain demonstrated a 50% higher EG-to-GA conversion titer with minimal genetic modifications compared to recently reported *E. coli* strain expressing *gox0313-aldA* genes which resulted only up to 4 g/L GA from 10 g/L EG (Yan et al., 2024).

**Fig. 3.**
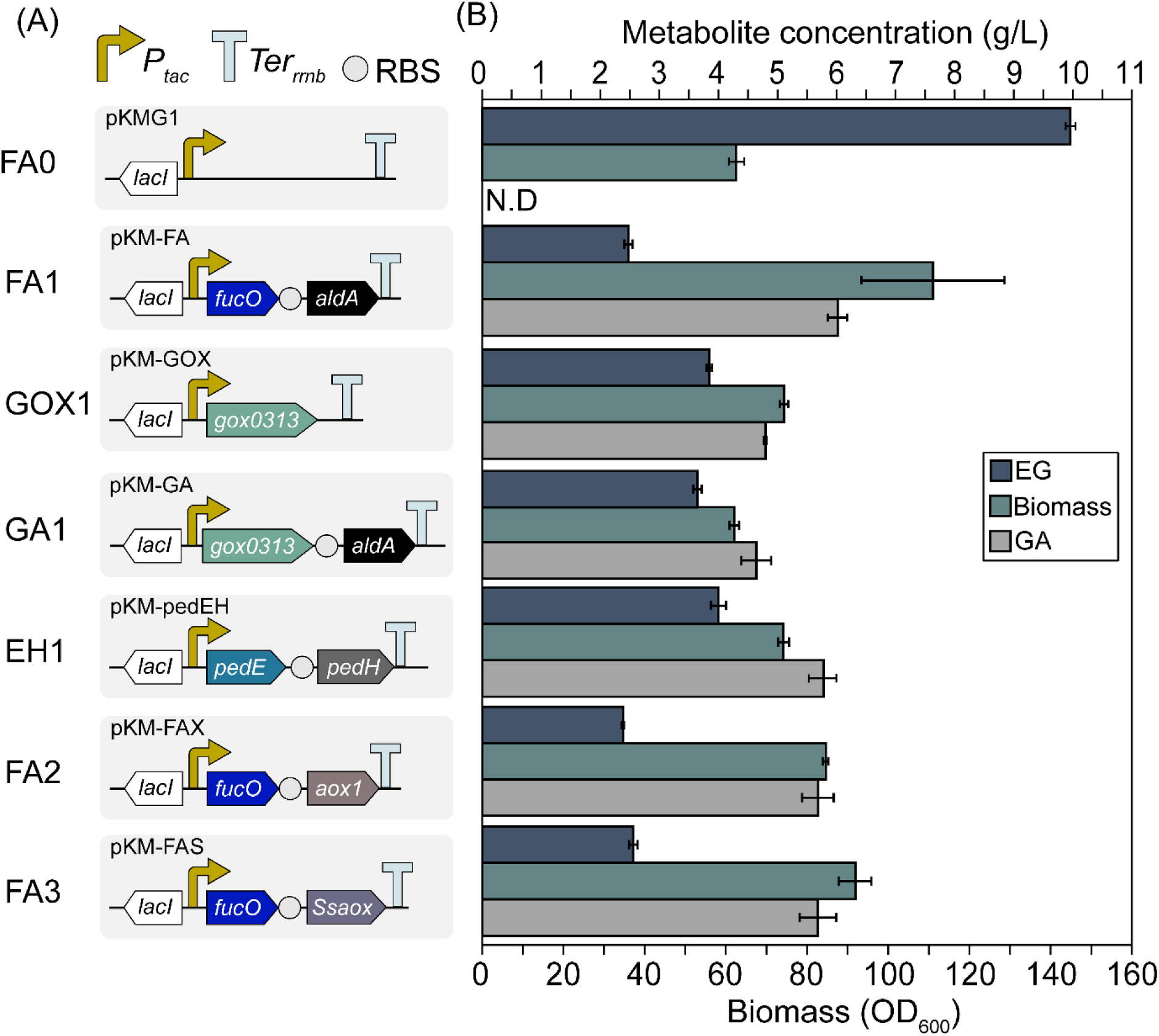
Construction scheme of the EG-to-GA biotransformation using *C. glutamicum*. (A) Schematic representation of the constructed plasmids used to produce GA. Symbols and abbreviations are as follows: *P_tac_*, tac inducible promoter; RBS, ribosome binding site; *Ter_rrnb_*, terminator. The presented genes and their encoding proteins are as follows: *fucO*^I7L/L8V^, propanediol oxidoreductase from *E. coli*; *gox0313*, alcohol dehydrogenase from *G. oxydans*; *aldA,* glycolaldehyde dehydrogenase from *E. coli; pedE,* alcohol dehydrogenases from *P. putida*; *pedH*, alcohol dehydrogenases from *P. putida*; *Ssaox*, glycolaldehyde dehydrogenase from *S. stipites*; and *aox1*, glycolaldehyde dehydrogenase from *P. pastoris*. (B) Biomass and GA titer obtained from flask cultivations of *C. glutamicum* strains harboring different plasmids in the presence of CGXII minimal media supplemented with 10 g/L EG and 40 g/L glucose for 48 h. Error bars represent the mean ± SD from triplicate. N.D = not detected.

As indicated in a previous report (Balola et al., 2024), the conversion of glycolaldehyde to GA is a bottleneck in this pathway. Therefore, to confirm if the conversion of glycolaldehyde to GA is also limiting for *C. glutamicum*, we tested two additional heterologous glycolaldehyde dehydrogenase from *Scheffersomyces stipitis* (encoded by *Ssaox*) and *Pichia pastoris* (encoded by *aox1*) which have shown good catalytic activity for the oxidation of aldehydes, to potentially improve the overall EG-to-GA conversion titer (Peña et al., 2018; Senatore et al., 2024). To do this, *aldA* gene in pKM-FA plasmid was replaced by *aox1* and *Ssaox*, resulting in the pKM-FAX and pKM-FAS plasmids. These plasmids were then separately transformed into *C. glutamicum* to generate FA2 and FA3 strains, respectively. After 48 h cultivation in shake flasks using media containing 10 g/L EG and 40 g/L glucose, the final GA titer obtained from the new strains were only 5.7 and 5.6 g/L, respectively, which did not exceed the titer obtained using the FA1 strain. This result suggested that glycolaldehyde-to-GA conversion might not be the rate-limiting reaction in this biosynthetic pathway. Therefore, the FA1 strain was selected for further optimization.

### 3.3 Optimization of gene expression level with promoter exchange

Engineering of promoter sequence is a feasible approach to balance gene expression and fine-tune metabolic pathways to enhance the targeted metabolic flux and subsequently product yields. So far, P_tac_ promoter was consistently used in all constructed plasmid for EG-to-GA conversion, resulting in a GA production of 6 g/L in FA1 strain. However, P_tac_ promoter is known to provide low-to-moderate expression level compared to constitutive promoters such as P*_tuf_* and P_H36_ (Ghiffary et al., 2022). To further enhance GA production, we swapped P*_tac_* promoter in pKM-FA with P*_tuf_* and P_H36_, resulting in pKTuf-FA and pKH-FA plasmid, respectively. These plasmids were then introduced into *C. glutamicum*, generating FA4 and FA5 strains, respectively. After 48 hours of cultivations, 10.6 and 9.4 g/L GA were obtained from FA4 and FA5 strains, respectively, representing a 77% and 56% increase compared to the FA1 strain (Fig. 4). Moreover, no remaining EG was detected at the end of FA4 cultivation, while around 2 g/L of EG was left in both FA1 and FA5 (Fig. 4), suggesting efficient EG-to-GA conversion of FA4 strain.

**Fig. 4.**
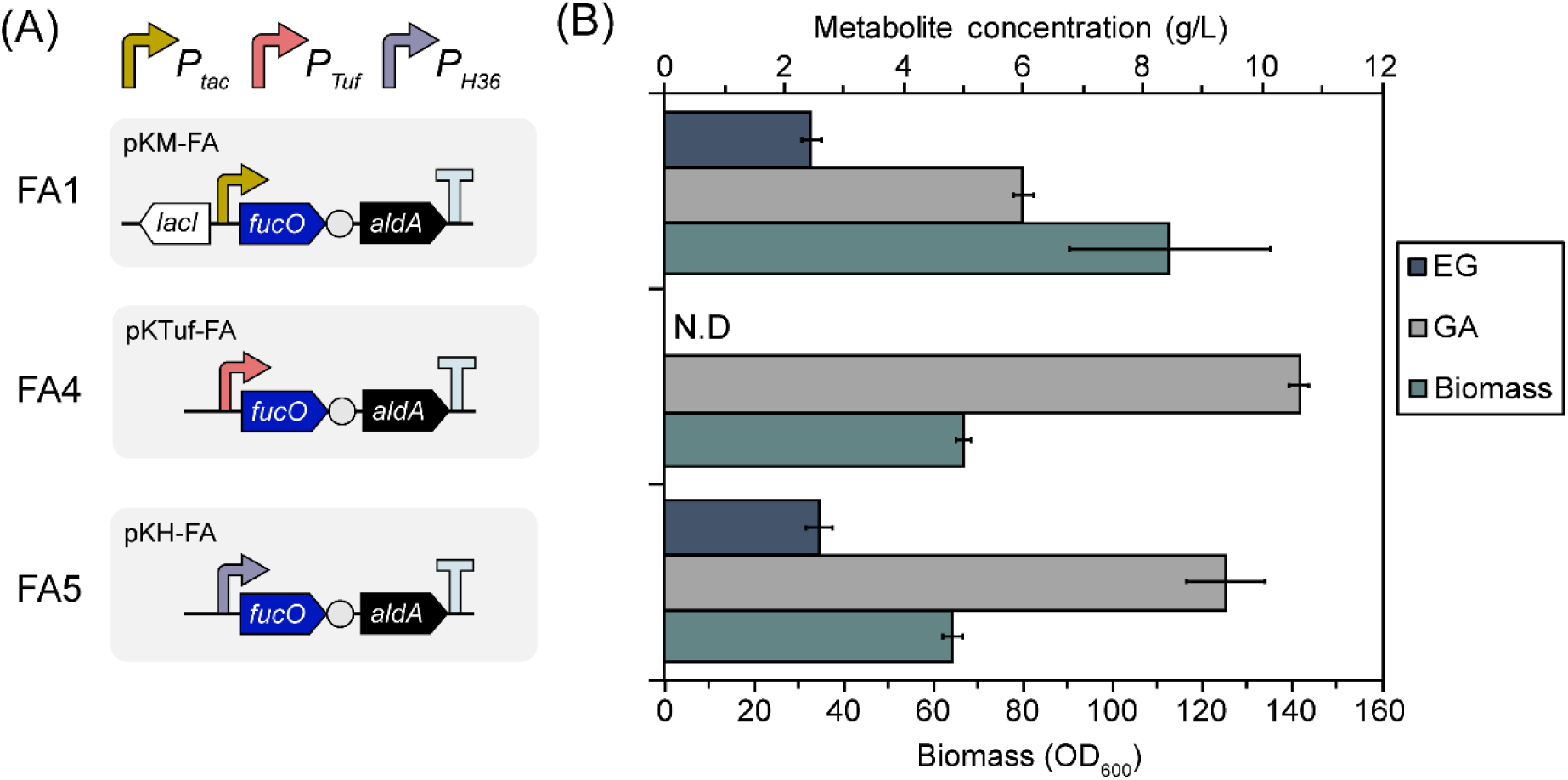
Effect of promoter on cell growth, EG utilization, and GA production. (A) Schematic representation of the constructed plasmids with different promoters used to produce GA. (B) Biomass, EG, and GA concentration obtained from flask cultivations of *C. glutamicum* strains harboring different plasmids in the presence of CGXII minimal media supplemented with 10 g/L EG and 40 g/L glucose for 48 h. Error bars represent the mean ± SD from triplicate. N.D = not detected.

This result is consistent with previous studies which have shown the following order of promoter strength in *C. glutamicum*: P_tac_ < P_H36_ *<* P*_tuf_* (Ghiffary et al., 2021). Although it is important to note that a stronger expression level does not always guarantee a higher titer of the target product due to the involvement of multiple cellular mechanisms. Of note, we observed a substantial decline in the final biomass of FA4 and FA5 compared to the FA1 strain (Fig. 4). This phenomenon is commonly found in engineered microbes designed to increase production of a target molecule because of a trade-off between maximizing growth and maximizing production, as the allocation of resources to one can negatively impact the other. This trade-off can be attributed to limited cellular resources and competition for metabolic precursors, among other factors (Sharma et al., 2024). Finally, based on GA production capability of the strains in the presence of glucose and EG media, FA4 with 10.6g/L EG production and minimal residual EG post cultivation was selected.

### 3.4 Two-stage biotransformation of EG to GA using repetitive resting cells of C. glutamicum

As the wild-type *C. glutamicum* requires glucose as a carbon and energy source for growth, a two-stage biotransformation strategy was explored. This approach consisted of an initial growth phase in CGXII minimal media with 40 g/L glucose to obtain sufficient biomass, followed by a production phase where the cells were transitioned to a resting state in minimal media containing 40 g/L EG as the sole carbon source for GA production. This strategy aimed to decouple cell growth from GA production that would potentially enhance the overall efficiency of the biotransformation process by directing cellular resources towards product biosynthesis.

To do this, the FA4 strain, which previously demonstrated the highest GA production in single-phase biotransformation, was cultivated in CGXII minimal media with 40 g/L glucose in shake flasks for 24 hours until it reached a stationary phase (OD_600_ ≈ 60). Subsequently, the biomass was harvested by centrifugation, resuspended in fresh CGXII minimal media containing 40 g/L EG as the sole carbon source, and incubated for 24 h at 30°C with shaking at 200 rpm. This cycle was then repeated every 24 h for an additional 2 days, and samples were taken every 24 h to measure EG and GA concentration as well as biomass (OD_600_).

The time-course profile of GA production using resting cells of the FA4 strain is shown in Fig. 5. The GA production titer ranged from around 26 to 38 g/L in each of biotransformation cycle, resulting in a cumulative GA titer, yield, and productivity of 98.8 g/L, 0.8 (gram GA/gram EG), and 1.37 g/L/h, respectively, over the 72-hours production phase. Notably, the OD_600_ of the resting cell culture was slightly decreased in each cycle throughout the total of 96-hour experiment, likely due to the cell loss during biomass harvesting. Nonetheless, to the best of our knowledge, the obtained cumulative GA titer is the highest reported to-date for similar biotransformation processes (Table 2).

**Fig. 5.**
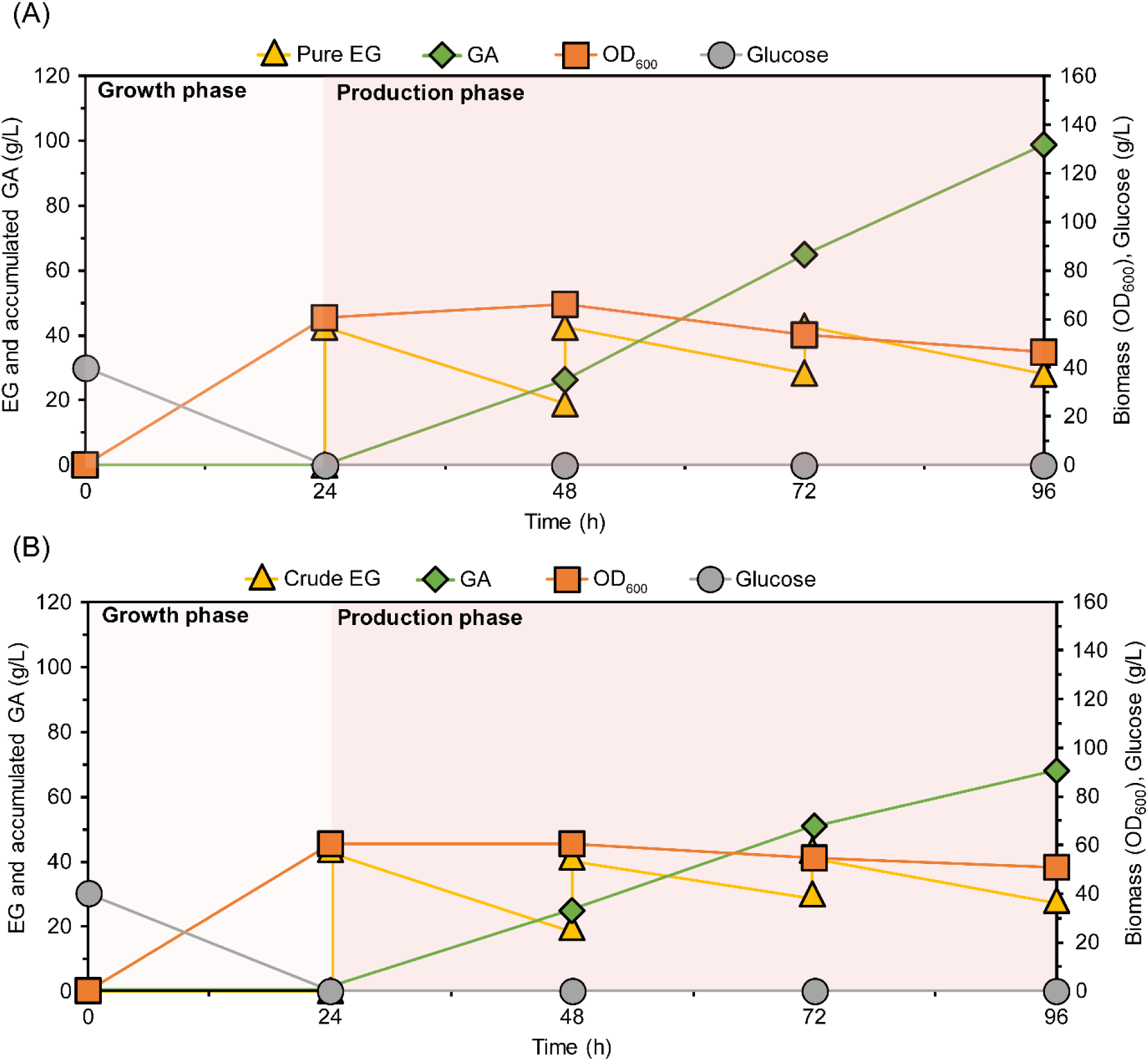
Schematic of two-stage biotransformation of EG to GA using repetitive resting cells of *C. glutamicum*. (A) Biomass growth, EG consumption, and GA production profile of *C. glutamicum* FA4 grown in minimal media during growth phase and supplemented with pure EG during production phase. (A) Biomass growth, EG consumption, and GA production profile of C. glutamicum FA4 grown in minimal media during growth phase, and supplemented with mock EG mixture (containing EG, TPA and PET) during production phase. Results are given as the average of n = 3. Detailed numbers are given in Table S4 and S5.

**Table 2.**
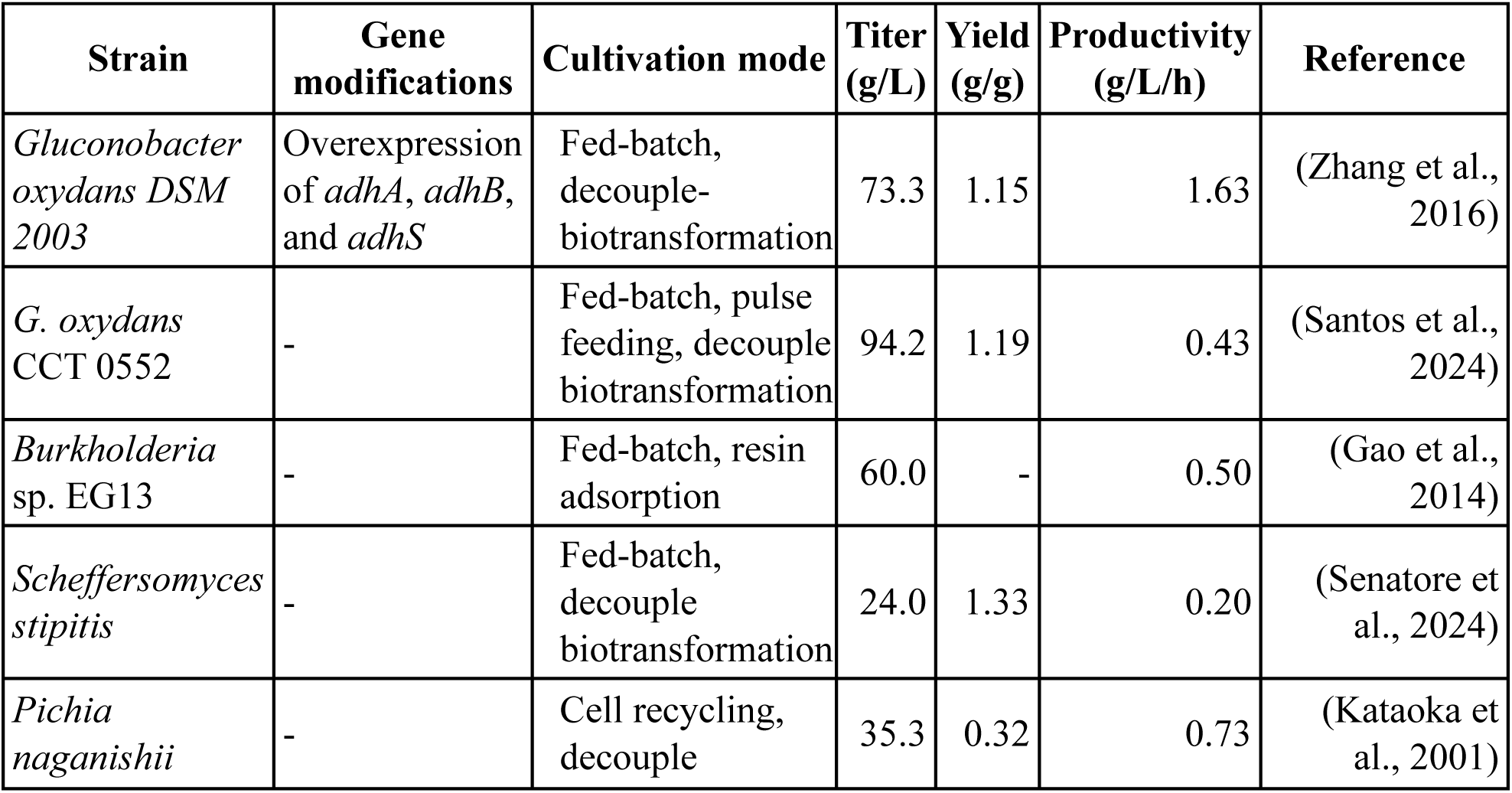
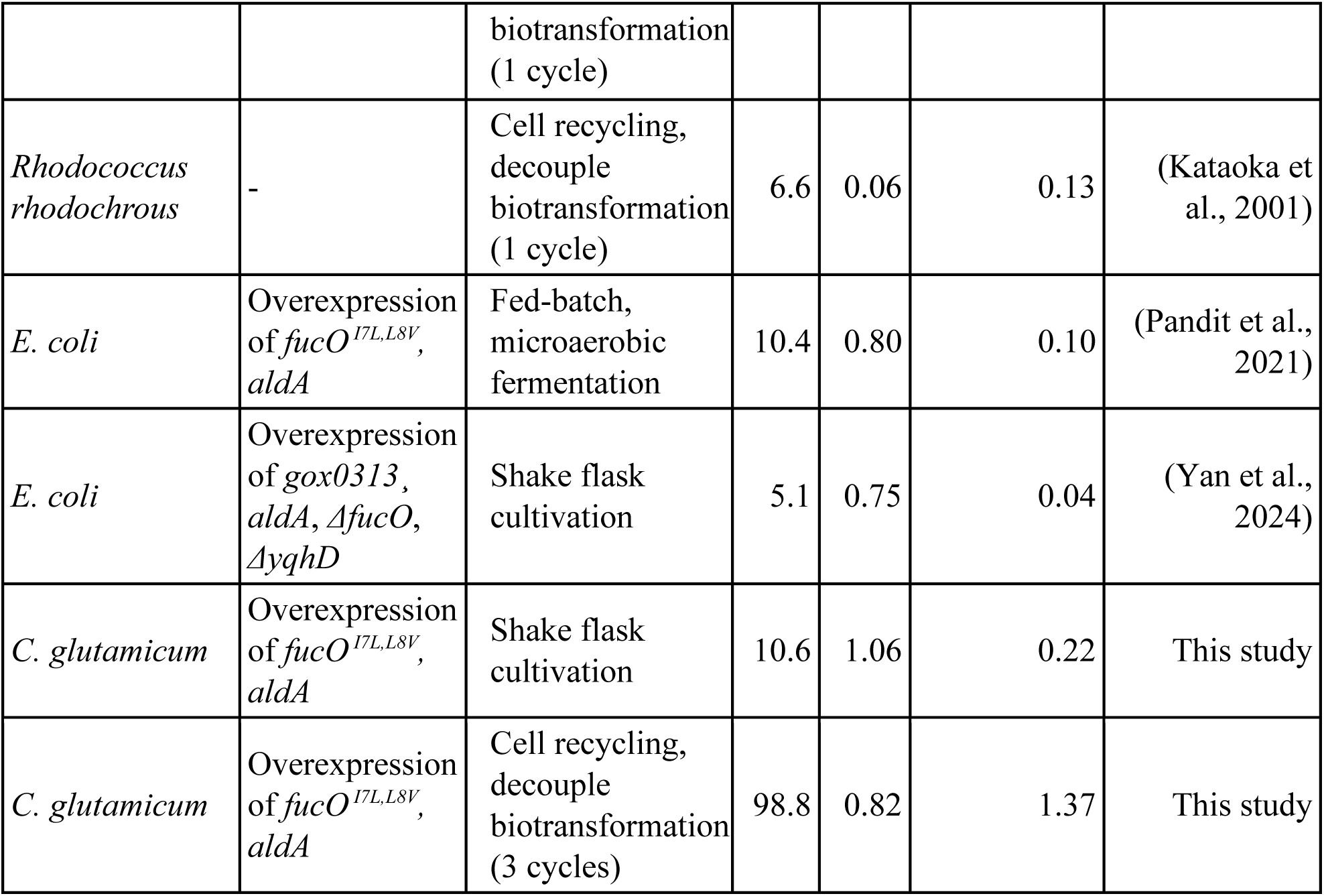
Recent work on GA production from EG through microbial biotransformation.

Additionally, we explored the potential of our developed biotransformation platform for the valorization of PET-plastic depolymerization products. Biological recycling of PET typically generates a mixture of its constituent monomers, EG and terephthalic acid (TPA), along with potentially unreacted PET oligomers or fragments. While TPA purification from the post PET-plastic degradation solutions can be achieved relatively easily, purification of EG is more challenging due to its high solubility in aqueous solution. For this initial feasibility study, we aimed to evaluate bioconversion of the remaining EG mixture derived under conditions where the PET depolymerization generated TPA monomer have been largely removed. Therefore, to simulate a relevant but simplified challenge for raw EG-to-GA bioconversion, we prepared a mock EG mixture containing 40 g/L EG, 2.5 g/L terephthalic acid (TPA), and 1 g/L PET as the biotransformation medium, supplemented with essential minerals as in CGXII minimal media but without glucose. The concentration of EG was chosen based on the previously optimized conditions, while the TPA and PET concentration were set to represent a significant byproduct of a post-PET degradation and to mimic the presence of undigested plastic fragments that might be present in a real-world scenario.

Using previously described two-stage biotransformation conditions, the FA4 strain was capable of converting this mock EG mixture to GA, with the GA titer varied from 17 to 26 g/L in each cycle and a cumulative GA production of approximately 67.3 g/L over the 72-hour biotransformation (Fig. 5). While this yield was lower than that obtained with pure EG under optimized conditions, it still demonstrates the feasibility of utilizing a complex feedstock derived from PET waste for GA production. Interestingly, the presence of PET fragments did not appear to significantly inhibit the biotransformation process, as the biomass (OD_600_) remained stable over 96-hour of biotransformation. However, the lower GA yield in the presence of TPA suggests a potential inhibitory effect of TPA on the enzymatic reaction or cell physiology. Further investigation into the specific mechanisms of this inhibition would be valuable.

## 4. Conclusions

In conclusion, we have developed a biotransformation platform for the production of GA from EG using the industrial workhorse *C. glutamicum*. We demonstrated that wild-type *C. glutamicum* exhibits tolerance to high concentrations of both EG and GA but cannot utilize these compounds as sole carbon sources. Through the heterologous expression of a synthetic EG oxidation pathway, we engineered *C. glutamicum* strains capable of converting EG to GA and further optimized the gene expression by promoter exchange which significantly enhanced GA production to 10.6 g/L in the final optimized strain, FA4. Moreover, the implementation of a two-stage biotransformation strategy utilizing repetitive resting cells of the FA4 strain resulted in a cumulative GA titer of 98.8 g/L in a 72-h production, representing a substantial improvement over single-phase cultivation and surpassing previously reported yields and productivity, reaching 0.8 (gram GA/gram EG), and 1.37 g/L/h, respectively. Also, by utilizing a mock EG mixture solution containing EG, terephthalic acid (TPA), and PET, we showcased a successful bioconversion of EG to GA in this complex medium, albeit lower titer than that obtained with pure EG. Overall, this work establishes *C. glutamicum* as a robust chassis for EG-to-GA bioconversion and provides a foundation for future research focused on further optimizing the process and implementing similar system for the sustainable production of chemicals from a waste-derived feedstock.

## Author contributions

**Mohammad Rifqi Ghiffary:** Conceptualization, Methodology, Investigation, Validation, Formal analysis, Data curation, Writing-original draft, Writing-reviewing & editing, Figure drawing. **Fong Tian Wong:** Conceptualization, Resources, Funding acquisition, Supervision, Project administration, Writing-review & editing. **Yee Hwee Lim:** Conceptualization, Resources, Funding acquisition, Supervision, Project administration, Writing-review & editing.

## Acknowledgements

We thank Ying Sin Koo (A*STAR ISCE^2^) for kindly providing the mock PET-degraded solution. This work is supported by the Agency for Science, Technology and Research (A*STAR) under C233017003, C233017004 and the Singapore Integrative Biosystems and Engineering Research Strategic Research & Translational Thrust (SIBER SRTT).

## Data Availability Statement

Data will be made available on request

