## Supplementary Information for "Production of Glycolic acid through Whole-Cell Bioconversion from PET Monomer Ethylene Glycol Using Engineered *Corynebacterium glutamicum*"

*Corresponding authors

Phone numbers: +65-XXXX-XXXX

### Supplementary Tables

#### Table S1

Primers and their DNA template used to construct plasmids in this study.

| Primer | Sequence (5’ → 3’) | DNA template | Final plasmid | Cloning method |
|---|---|---|---|---|
| P1 | gtgagcggataacaatttcacacaggaaacagaattaaaagatgatggctaacagaatgctggtgaac | gDNA of *E. coli* K-12 | pKM-FA | Gibson assembly |
| P2 | ttaccaggcggtatggtaaagctctac | | | |
| P3 | ggatattgtagagctttaccataccgcctggtaagaaaggagaggattgcatgtcagtacccgttcaacatcctatg | gDNA of *E. coli* K-12 | | |
| P4 | gcatgcaagcttggcgtaatcatggtcttaagactgtaaataaaccacctgggtctg | | | |
| P5 | gaccatgattacgccaagcttgc | pKMG1 plasmid | | |
| P6 | cttttaattctgtttcctgtgtgaaattgttatccg | | | |
| P7 | gtgagcggataacaatttcacacaggaaacagaattaaaagatgacaataagatcgctacccgcc | gDNA of *P. putida* | pKM-pedEH | Gibson assembly |
| P8 | cgacgcggggatcgggtcatgcaatcctctcctttctcagcgctgcgcagtcttg | | | |
| P9 | atgacccgatccccgcgt | gDNA of *P. putida* | | |
| P10 | gcatgcaagcttggcgtaatcatggtcttacggcttggcgcttgcc | | | |
| P5 | gaccatgattacgccaagcttgc | pKMG1 plasmid | | |
| P6 | cttttaattctgtttcctgtgtgaaattgttatccg | | | |
| P11 | gagcggataacaatttcacacaggaaacagaattaaaagatggctgatacaatgctcgcc | gDNA of *G. oxydans* | pKM-GOX | Gibson assembly |
| P12 | gcatgcaagcttggcgtaatcatggtctcaggaccggaagtcgagcac | | | |

| | | | | |
|---|---|---|---|---|
| P5 | gaccatgattacgccaagcttgc | pKMG1 plasmid | | |
| P6 | cttttaattctgtttcctgtgtgaaattgttatccg | | | |
| P11 | gagcggataacaatttcacacaggaaacagaattaaaagatggctgatacaatgctcgcc | gDNA of *G. oxydans* | pKM-GA | Gibson assembly |
| P13 | tcaggaccggaagtcgagcac | | | |
| P14 | ccggtcctgagaaaggagaggattgcatgtcagtacccgttcaacatcc | gDNA of *G. oxydans* | | |
| P15 | gcatgcaagcttggcgtaatcatggtcttaagactgtaaataaaccacctgggtctg | | | |
| P5 | gaccatgattacgccaagcttgc | pKMG1 plasmid | | |
| P6 | cttttaattctgtttcctgtgtgaaattgttatccg | | | |
| P1 | gtgagcggataacaatttcacacaggaaacagaattaaaagatgatggctaacagaatgctggtgaac | gDNA of *E. coli* K-12 | pKM-FAS | Gibson assembly |
| P2 | ttaccaggcggtatggtaaagctctac | | | |
| P16 | gatattgtagagctttaccataccgcctggtaagaaaggagaggattgcatgtccaaagtatttatcacgggtgcaac | Synthetic *Ssaox* gene | | |
| P17 | catgcaagcttggcgtaatcatggtctcaattgcgcttaacgagatagtcaatg | | | |
| P5 | gaccatgattacgccaagcttgc | pKMG1 plasmid | | |
| P6 | cttttaattctgtttcctgtgtgaaattgttatccg | | | |
| P1 | gtgagcggataacaatttcacacaggaaacagaattaaaagatgatggctaacagaatgctggtgaac | gDNA of *E. coli* K-12 | pKM-FAX | Gibson assembly |
| P2 | ttaccaggcggtatggtaaagctctac | | | |
| P18 | gatattgtagagctttaccataccgcctggtaagaaaggagaggattgcatggcaatcccagaagagtttgacatc | Synthetic *aox1* gene | | |
| P19 | gcatgcaagcttggcgtaatcatggtcctaaaagcgcgccaagccc | | | |
| P5 | gaccatgattacgccaagcttgc | pKMG1 plasmid | | |
| P6 | cttttaattctgtttcctgtgtgaaattgttatccg | | | |
| P20 | gccaccacgaagtccaggaggacatacaatgatggctaacagaatgctggtgaac | gDNA of *E. coli* K-12 | pKTuf-FA | Gibson assembly |
| P2 | ttaccaggcggtatggtaaagctctac | | | |

| | | | | |
|---|---|---|---|---|
| P3 | ggatattgtagagctttaccataccgcctggtaagaaaggagaggattgcatgtcagtacccgttcaacatcctatg | gDNA of *E. coli* K-12 | | |
| P4 | gcatgcaagcttggcgtaatcatggtcttaagactgtaaataaaccacctgggtctg | | | |
| P5 | gaccatgattacgccaagcttgc | pKTG1 plasmid | | |
| P21 | tgtatgtcctcctggacttcgtggtg | | | |
| P22 | ccttggttggtaggagtagcatgggatccatgatggctaacagaatgctggtgaac | gDNA of *E. coli* K-12 | pKH-FA | Gibson assembly |
| P2 | ttaccaggcggtatggtaaagctctac | | | |
| P3 | ggatattgtagagctttaccataccgcctggtaagaaaggagaggattgcatgtcagtacccgttcaacatcctatg | gDNA of *E. coli* K-12 | | |
| P4 | gcatgcaagcttggcgtaatcatggtcttaagactgtaaataaaccacctgggtctg | | | |
| P5 | gaccatgattacgccaagcttgc | pKHG1 plasmid | | |
| P23 | ggatcccatgctactcctaccaac | | | |

**Table S2.** DNA sequence of the codon-optimized gene used in this study.

| Gene name | Sequence (5' → 3') |
|---|---|
| Codon-optimized *Ssaox* | ATGTCCAAAGTATTTATCACGGGTGCAACTGGCTATATTGGAGGA<br>CAGGTTCTTTATGAGCTGCTTAATAACAAGGATGGACGGAAATAT<br>GACGTGACCGCGCTGGTACGCTCTCAACAAAGGCCGAGAAGCT<br>TCTTATTGCGACGAATAATCAAATTTCAACGGTTATCGGTTCACTT<br>GATGATGTAGACTTTATCAAGCAACAAGTAGAAGCTAATGATATT<br>ATCATTAATACGGCCAACGTAGACCATGTTCCATCTGCTCAAGCT<br>GTTTCAGACGCTCTCGTGGCGTCGAAGGACAAGAAGATTTATATT<br>CATACGTCAGGCACTTCAATTCTTGGCGACGGCCTCTCGCCAGAC<br>AAAGGAGACTCTCACAAGGTTTACTCGGATAAATATTCCATTGAT<br>GAGATTAATTCTTTTCCAGACACCCAGCCGCACCGCCCCGTAGAC<br>CGTATTGTACTTGACATCCACAATAAAAACCCAAATGTACAAGTG<br>GTAGTCATTTGTCCGTCTACTATTTACGGAATCTCAAACGGTTACG<br>ATAATCTTGTATCGGCCCAGGTTCCATTGCTCATCACTTCCTTTGT<br>TAAATATGGTAAAGGATACACCGTGTATAAGGGTGACGAGATTTG<br>GAACCATATCCATATCAAGGACTTGGGTGACTTGTATTATCTCATT<br>CTGACTAAGCTGCAATCCGGCGAGGATATTCCCGTCAACGGAACC<br>GGTTATTACTTCGGCAGCTTGGCAATTGAAGGAGAGGAGACCATT<br>TCAAACGAGCCGTCGTCGATCGAGCACCGGTGGCGCCAGGTAAG<br>CGAGGTTGTGGCTGAAAAGCTCTTTTCGAAAGGCCTCATTGGATC<br>AAAGGAAGTTGTCTCTCTTGAACCGCAAGAAATTGCCAAGATCAA<br>CGATTCGGAATGGTCACCATTTTACTGGGGTACCTACTCGCGCTC<br>ACGTGGAGATAACGGCTATGCCATTGGTTGGAAGCCAAAGTTCAC<br>TAGCAACAAGGAGTTCTTCGATTCTTTTGATGCGGACATTGACTA<br>TCTCGTTAAGCGCAATTGA |
| Codon-optimized *aox1* | ATGGCAATCCCAGAAGAGTTTGACATCTTGGTACTCGGCGGAGGA<br>TCTAGCGGCAGCTGTATCGCCGGACGCCTTGCTAATCTGGATCAT<br>TCCCTCAAGGTAGGATTGATTGAGGCAGGAGAGAACAACCTTAA<br>CAACCCATGGGTCTATCTGCCAGGAATTTACCCACGCAACATGAA<br>ACTCGACTCTAAAACGGCTTCGTTCTATACGTCAAATCCCTCACCC<br>CATCTCAACGGTCGCCGCGCGATTGTTCCGTGTGCCAATGTTTTGG<br>GAGGTGGCTCTAGCATCAACTTTATGATGTACACGCGGGGATCCG<br>CATCTGACTATGATGACTTTCAGGCTGAGGGTTGGAAAACCAAAG<br>ACCTCCTCCCATTGATGAAAAAAACCGAGACGTATCAGCGTGCCT<br>GCAATAACCCTGATATTCATGGCTTTGAGGGACCAATTAAAGTAT<br>CCTTCGGAAACTACACCTACCCCGTATGCCAGGACTTTCTTCGCG<br>CAAGCGAGTCACAGGGAATTCCTTATGTGGACGATTTGGAAGATT<br>TGGTCACGGCTCATGGTGCGGAACACTGGTTGAAATGGATCAACC<br>GGGATACTGGCCGGCGCAGCGACTCAGCACACGCATTCGTACATT<br>CGACTATGCGTAACCATGACAACTTGTATCTCATCTGTAACACCA<br>AAGTTGACAAAATCATTGTTGAGGATGGACGTGCTGCAGCTGTTC<br>GTACGGTGCCGTCAAAGCCTCTTAACCCAAAAAAGCCGTCACACA<br>AAATTTATCGTGCTCGGAAGCAAATTGTACTCTCGTGCGGAACGA<br>TTTCATCGCCTCTGGTTCTGCAGCGTTCAGGTTTTGGTGATCCTAT<br>TAAGTTGCGGGCGGCCGGAGTTAAACCCCTCGTGAATCTTCCTGG<br>TGTAGGACGTAATTTCCAGGATCATTATTGTTTTTCTCACCCTAT |

| | |
|---|---|
| | CGTATCAAGCCTCAGTATGAATCGTTTGACGATTTTGTCCGTGGTG<br>ACGCTGAAATCCAAAAACGCGTATTCGACCAATGGTACGCAAAT<br>GGTACCGGCCCATTGGCAACCAACGGCATTGAGGCAGGTGTAAA<br>GATCCGGCCAACTCCAGAAGAACTCTCTCAAATGGATGAATCATT<br>CCAAGAAGGATACCGCGAATATTTCGAGGATAAACCAGACAAAC<br>CCGTAATGCACTACAGCATCATCGCTGGCTTTTTCGGTGACCACA<br>CTAAAATTCCGCCAGGAAAGTACATGACTATGTTCCACTTCCTCG<br>AGTACCCATTTTCTCGGGGATCCATCCACATTACGTCACCGGATC<br>CCTATGCGGCACCTGATTTTGATCCCGGATTTATGAATGACGAAC<br>GCGACATGGCACCTATGGTGTGGGCCTACAAGAAAAGCCGTGAA<br>ACCGCCCGGCGCATGGATCACTTCGCAGGCGAGGTAACTTCACAT<br>CACCCACTCTTCCCATACTCGTCAGAAGCGCGTGCCCTGGAGATG<br>GATCTCGAAACTTCGAATGCGTATGGAGGACCCCTTAATCTGTCA<br>GCCGGACTGGCGCACGGTAGCTGGACCCAACCATTGAAGAAGCC<br>AACGGCAAAGAACGAGGGCCATGTAACGTCAAACCAGGTTGAAC<br>TGCATCCTGACATTGAATATGACGAAGAGGATGACAAGGCGATT<br>GAGAACTACATTCGTGAGCATACCGAAACTACTTGGCATTGCCTG<br>GGAACCTGTTCAATCGGTCCCCGTGAGGGATCTAAGATTGTAAAA<br>TGGGGCGGAGTGTTGGACCATCGGTCGAATGTCTACGGAGTGAA<br>GGGCTTGAAGGTGGGAGATTTGAGCGTTTGCCCAGACAATGTTGG<br>CTGTAACACCTATACTACCGCATTGCTGATCGGAGAAAAAACCGC<br>GACCCTGGTGGGTGAGGATCTTGGTTATTCAGGAGAGGCATTGGA<br>CATGACCGTGCCGCAATTCAAGCTGGGAACGTACGAAAAGACGG<br>GCTTGGCGCGCTTTTAG |

**Table S3.** Remaining EG and glucose, GA titer, and biomass ($OD_{600}$) from flask cultivation of the *C. glutamicum* strains constructed in this study.

| Strain | Remaining EG (g/L) | Remaining glucose (g/L) | GA titer (g/L) | Biomass ($OD_{600}$) |
|---|---|---|---|---|
| FA0 | 10 ± 0.05 | N.D | N.D | 62.21 ± 1.21 |
| FA1 | 2.45 ± 0.15 | N.D | 6.00 ± 0.60 | 112.61 ± 22.52 |
| FA2 | 2.46 ± 0.09 | N.D | 5.67 ± 0.27 | 82.64 ± 0.60 |
| FA3 | 2.49 ± 0.14 | N.D | 5.58 ± 0.31 | 91.82 ± 4.00 |
| FA4 | N.D | N.D | 10.63 ± 0.16 | 66.57 ± 1.86 |
| FA5 | 2.58 ± 0.22 | N.D | 9.39 ± 0.66 | 64.02 ± 2.16 |
| GOX1 | 3.82 ± 0.06 | N.D | 4.78 ± 0.02 | 74.30 ± 1.01 |
| GA1 | 3.59 ± 0.22 | N.D | 4.63 ± 0.25 | 62.04 ± 1.15 |
| EH1 | 4.03 ± 0.52 | N.D | 5.81 ± 0.26 | 75.99 ± 0.86 |

* Error bars represent the mean and the standard deviation from triplicate experiments.

* N.D = Not detected

**Table S4.** Remaining EG and glucose, GA titer, and biomass ($OD_{600}$) from two-stage biotransformation of pure EG to GA using repetitive resting cells of *C. glutamicum* FA4 strain.

| Time | Remaining EG (g/L) | Remaining glucose (g/L) | GA titer (g/L) | Biomass ($OD_{600}$) |
|---|---|---|---|---|
| 0 | N.D | 40.00 ± 0.00 | N.D | 2.45 ± 0.04 |
| 24 | N.D | N.D | N.D | 60.44 ± 6.46 |
| 25 | 42.64 ± 1.54 | N.D | N.D | 60.44 ± 6.46 |
| 48 | 19.01 ± 1.20 | N.D | 26.39 ± 1.01 | 65.98 ± 15.35 |
| 49 | 42.62 ± 2.21 | N.D | N.D | 65.98 ± 15.35 |
| 72 | 28.45 ± 3.77 | N.D | 38.43 ± 8.13 | 53.43 ± 1.48 |
| 73 | 42.79 ± 1.46 | N.D | N.D | 53.43 ± 1.48 |
| 96 | 28.11 ± 5.65 | N.D | 34.04 ± 3.52 | 46.63 ± 0.57 |

* Error bars represent the average and the standard deviation from triplicate experiments

**Table S5.** Remaining EG and glucose, GA titer, and biomass ($OD_{600}$) from two-stage biotransformation of raw EG to GA using repetitive resting cells of *C. glutamicum* FA4 strain.

| Time | Remaining EG (g/L) | Remaining glucose (g/L) | GA titer (g/L) | Biomass ($OD_{600}$) |
|---|---|---|---|---|
| 0 | N.D | 40.00 ± 0.00 | N.D | 2.33 ± 0.02 |
| 24 | N.D | N.D | N.D | 60.44 ± 6.46 |
| 25 | 42.64 ± 1.54 | N.D | N.D | 60.44 ± 6.46 |
| 48 | 18.48 ± 0.34 | N.D | 23.92 ± 0.77 | 60.37 ± 5.82 |
| 49 | 40.27 ± 2.11 | N.D | N.D | 60.37 ± 5.82 |
| 72 | 28.89 ± 2.32 | N.D | 26.19 ± 3.00 | 54.86 ± 5.49 |
| 73 | 41.25 ± 1.12 | N.D | N.D | 54.86 ± 5.49 |
| 96 | 27.55 ± 1.33 | N.D | 17.22 ± 3.22 | 50.91 ± 15.17 |

* Error bars represent the average and the standard deviation from triplicate experiments

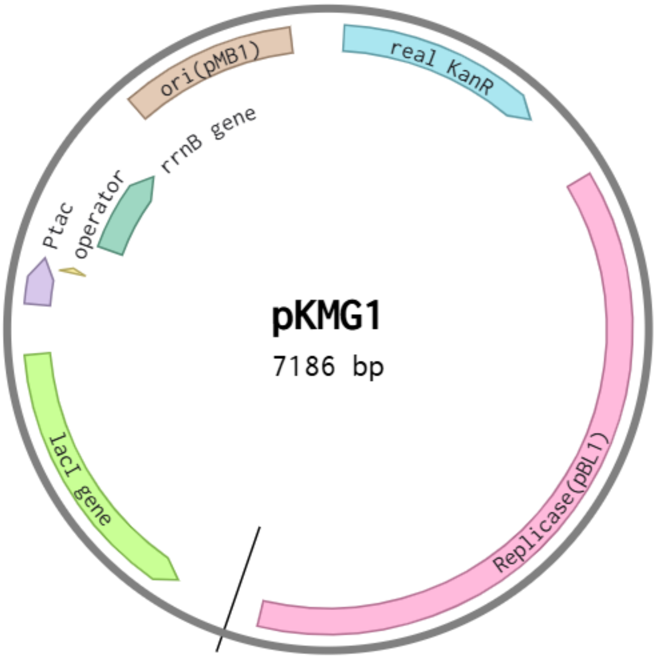


Figure S1. The map for pKMG1 plasmid

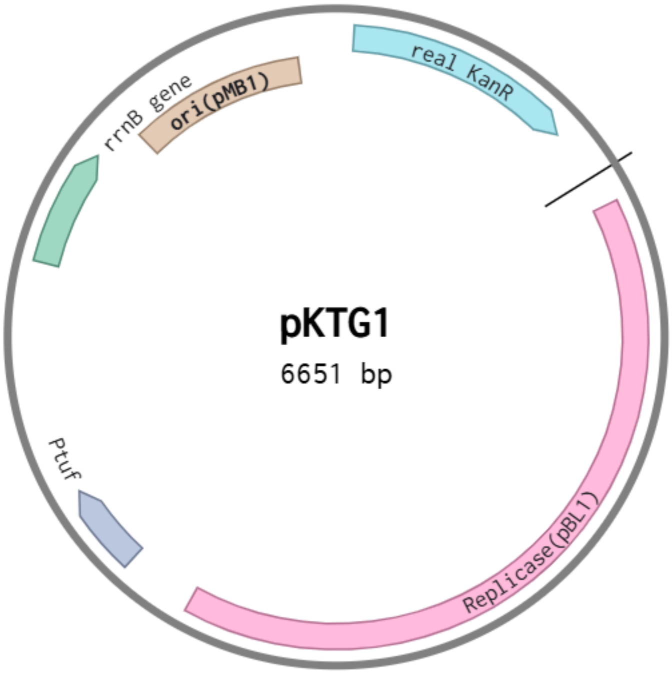


Figure S2. The map for pKTG1 plasmid

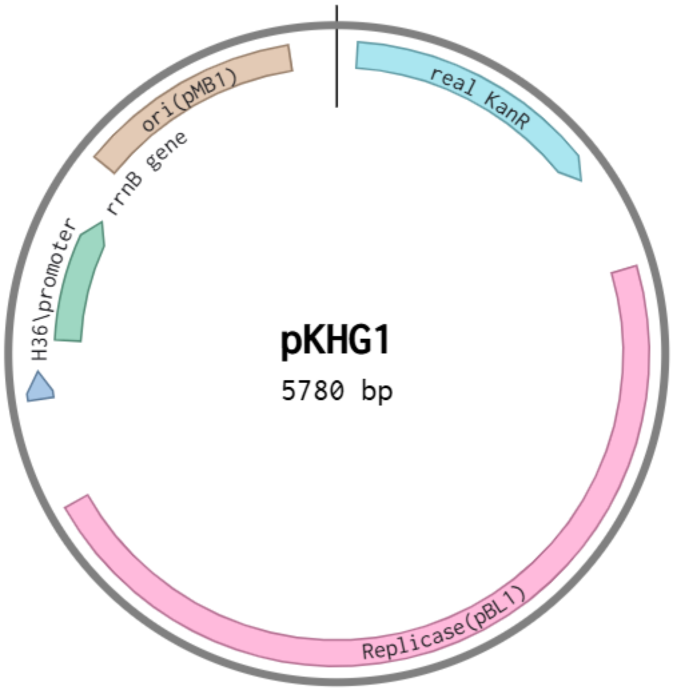


Figure S3. The map for pKHG1 plasmid

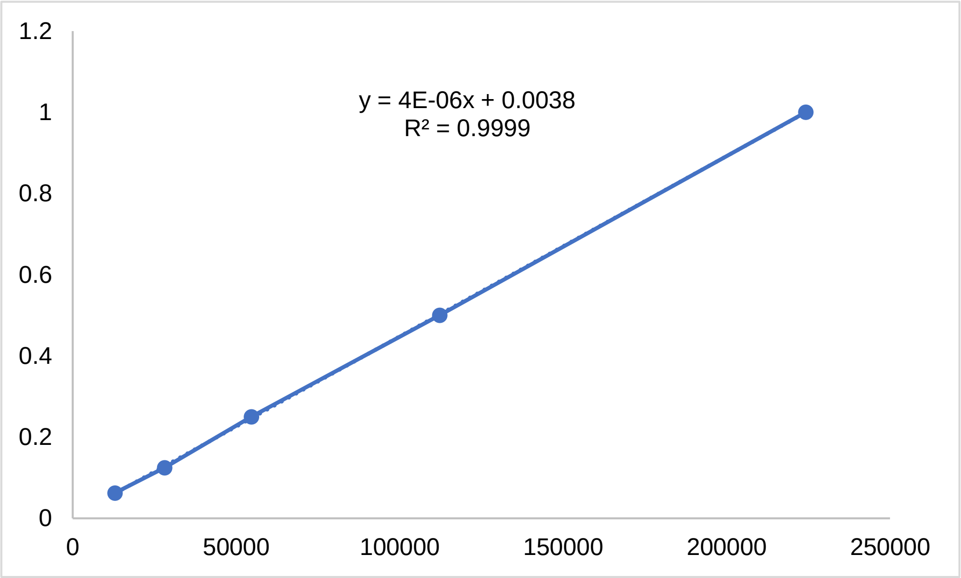


Figure S4. Calibration curve for EG

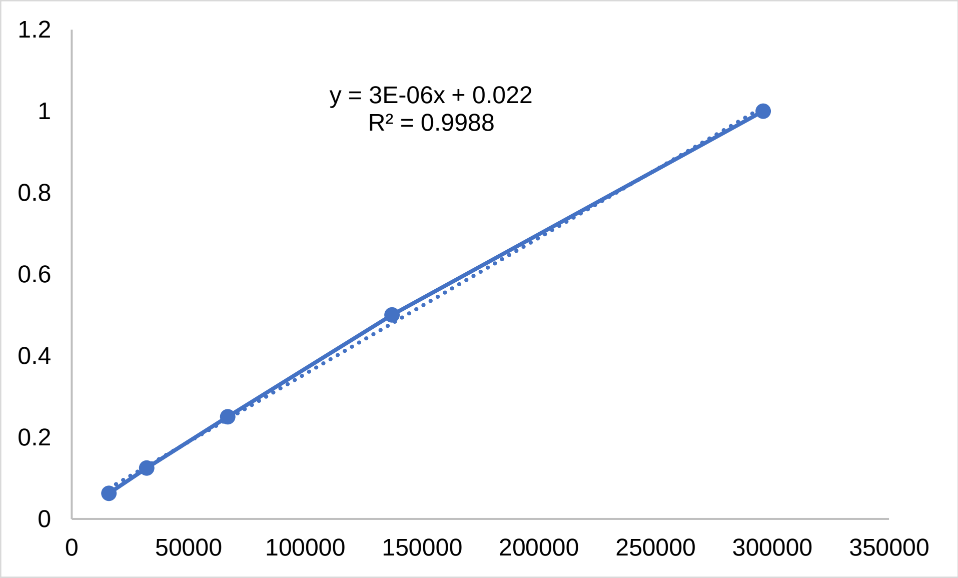


Figure S5. Calibration curve for glucose

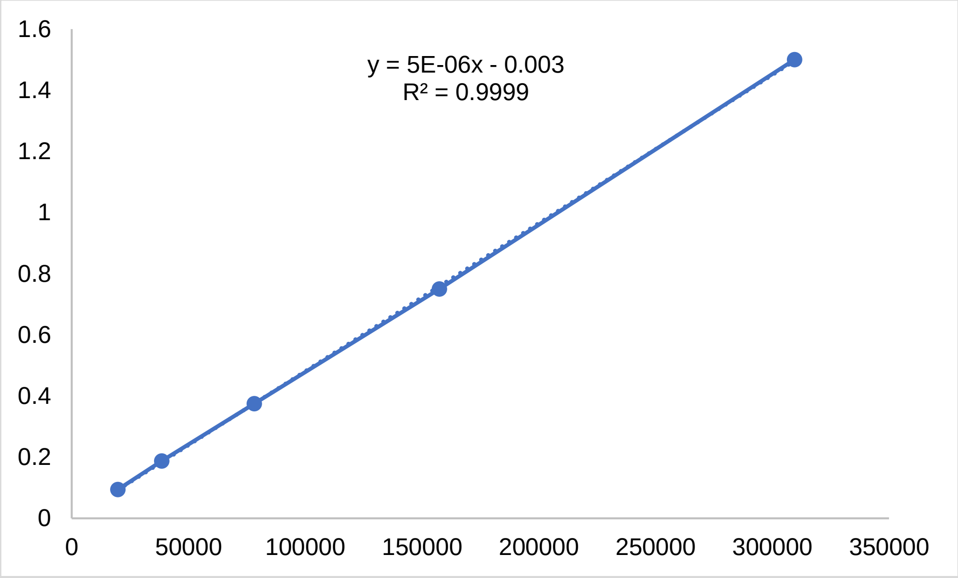


Figure S6. Calibration curve for GA
